# Transcriptional changes suggest a major involvement of Gibberellins in *Trifolium pratense* regrowth after mowing

**DOI:** 10.1101/775841

**Authors:** Denise Brigitte Herbert, Thomas Gross, Oliver Rupp, Annette Becker

## Abstract

Red clover (*Trifolium pratense*) is used worldwide as a fodder plant due its high nutritional value. In response to mowing, red clover exhibits specific morphological traits to compensate the loss of biomass. The morphological reaction is well described, but knowledge of the underlying molecular mechanisms are still lacking. Here we characterize the molecular genetic response to mowing of red clover by using comparative transcriptomics in greenhouse conditions and agriculturally used field. The analysis of mown and control plants revealed candidate genes possibly regulating crucial steps of the genetic network governing the regrowth reaction. In addition, multiple identified gibberellic acid (GA) related genes suggest a major role for GA in establishing the regrowth morphology of red clover. Mown red clover plants showing this regrowth morphology were partially “rescued” by exogenous GA application, demonstrating the influence of GA during regrowth. Our findings provide insights into the physiological and genetic processes of mowing red clover, to serve as a base for red clover yield improvement.

## Introduction

*Trifolium pratense* (red clover) is an important worldwide forage crop and thus of great economic interest. This perennial plant offers several advantages like a high protein content and soil improving characteristics, which can reduce the use of artificial nitrogen application and can enhance intake in livestock. Well-known disadvantages of red clover include poor persistence under several land use scenarios, like grazing or cutting [1–3]. *T. pratense* is a member of the Fabaceae (or legumes), which are, due to their economic value, among the most examined families in the plant kingdom with genome sequences available for species like *Medicago truncatula* (barrel clover) [4], *Lotus japonicus* (birdsfoot trefoil) [5], *Glycine max* (soy) [6], *Phaseolus vulgaris* (common bean) [7], *Cicer arietinum* (chickpea) [8], *Vigna unguiculata* (cowpea) [9],*Trifolium subterraneum* (subterranean clover) [10] *Trifolium medium* (zigzag clover) [11], and *T. pratense* (red clover) [12, 13].

Facing today’s challenges such as an increased demand on food production in an era of global climate change together with the aim to solve these problems in an environmental friendly and sustainable way requires improvement of forage crops like *T. pratense* [14, 15]. *T. pratense* breeding aims to offer genotypes with improved key agronomic traits (dry matter yield, high quality, resistance to diseases and abiotic/biotic stress, persistency, [16]), while improving its regrowth ability [2, 17]. Unfortunately, the morphological investigations of several *T. pratense* populations showed a correlation of persistency with non-favorable traits, like small plant size and prostrate growth habit [18]. Moreover, most *T. pratense* cultivars or accessions are locally adapted and require their specific local conditions to show the favored traits [19, 20], which decreases the stability for individual traits in breeding efforts [21]. *T. pratense* exhibits significant intraspecific variation due to high intrapopulation genetic diversity, thus, persistence and performance in response to mowing or cutting, depends on the variety, as well as developmental stage at the moment of damage [22–25].

Persistency can be defined as a sustained forage yield over several growing periods [26] and is a complex trait influenced by a variety of abiotic and biotic factors, and the regrowth ability of a plant [27]. Plants with high regrowth ability can survive more frequent and intense biomass loss and could be therefore more persistent. Decapitation or biomass loss due to herbivory or mowing triggers a complex reaction affected by environmental conditions, plant morphology, architecture, developmental stage and genotype [22]. After decapitation, the first stress response in other legumes like *Medicago sativa* and *Pisum sativum* involves the production of phytohormones: cytokinine, auxin, and strigolactones [28–30]. In addition, the mobilization of energy reserves is activated [31]. Phenotypic plasticity of plant architecture in combination with alterations of hormone concentrations can be observed in *P. sativum (pea)* and *T. pratense* after decapitation [25,30,32]. However, the molecular processes allowing plants to thrive even after an enormous loss of biomass remain still unclear, even in *Arabidopsis thaliana* [33, 34].

Here, we compare the transcriptomes of mown (cut) vs. unmown (uncut) *T. pratense* plants from two different field locations on the Biodiversity Exploratory “Heinich-Dün” [35] and greenhouse grown plants. Our field samples were subjected to standard agricultural treatment and we can thus discriminate transcriptional changes caused by abiotic factors and biotic interactions in the field from those regulating regrowth. We present the identification and *in silico* characterization of putative developmental regulators differentially expressed in the regrowth phase after mowing in the field and in the greenhouse that may contribute to the regrowth response of *T. pratense* and demonstrate that gibberellic acid (GA) is a major regulator of specific aspects of the regrowth morphology in red clover.

## Material and Methods

### Plant growth conditions, GA treatment, tissue sampling, and RNA extraction, cDNA library construction and RNA-Seq

Plant material for RNA-Seq was collected from three locations (fields and greenhouse, Fig. 1 A and table S1, thereby one field location includes two neighboring field sites). Field plant tissue for RNA-Seq was sampled on 11.06.2014 within the area of the Biodiversity Exploratory “Hainich-Dün” [35], located in Thuringia, Germany. Material was sampled on four neighboring sites; two mown pastures and two meadows that were not mown (FaM, FaNM, FbM, FbNM). For the greenhouse samples, seeds of regional *T. pratense* populations (from a region covering mainly Thuringia, Saxony, Saxony-Anhalt, Thuringian Forest and Uckermarck, Germany) were obtained from the Rieger Hofmann seed company (Blaufelden, Germany). Plants were grown in 23 °C with 16 h of light in pots of 12 cm diameter. Plants in the greenhouse were watered daily and compound fertilizer (8’8’6’+) was given every ten days. After 122 days after sowing, half of the plants were cut to 5 cm (GM and GNM). Material from mown plants was sampled approximately 14 days after mowing/cutting, to avoid sequencing of the transcripts related to the first stress response [36]. After collection, the samples were snap frozen in liquid nitrogen. For each site and the greenhouse two biological replicates of four pooled plants (shoot and leaf material) each were collected.

**Fig. 1:**
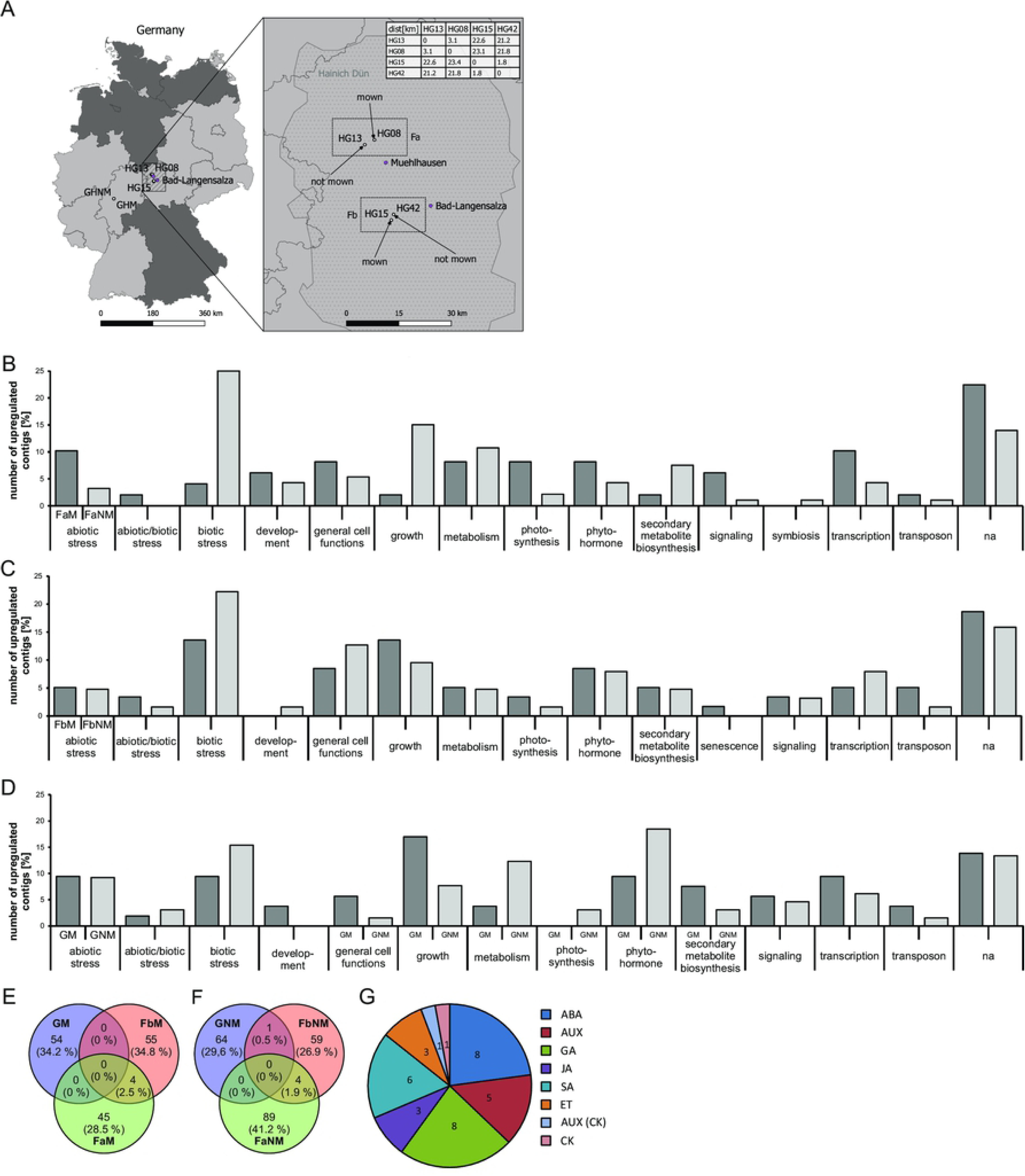
Overview of sampling locations and classification of DEGs. A: Overview of the sampling locations for the plant material. Names of the fields belonging to the Biodiversity Exploratory or greenhouse populations are shown, as well as the conditions (mown/cut and not mown/uncut, HG 15 and HG 42; HG13 and HG 08). Distances between sampling locations in the field have been estimated. B-D: Classification of DEGs with a |log2FoldChange| <2. Percentage share of each class to the corresponding gene list is shown in bar charts B: Classes of DEGs from field a mown vs. unmown. C: Classes of DE field b mown vs. not mown. D: Classes of DE contigs of mown plants grown in the greenhouse vs. unmown plants. E-F: Shared genes between the different treatments and locations. The Venn diagrams show the number of shared upregulated gene within the “mown” samples (E) and the number of shared genes within the “not mown” samples (F), blue circles indicate greenhouse data, green field a and red field b. G: Number of genes belonging to the class “phytohormones” within the DEG list of field (a and b) and greenhouse transcriptomes. The pie chart shows the number of the different plant phytohormones (absicic acid, ABA; auxin, AUX; genes common between the auxin and cytokinin pathway, AUX/CK; cytokinin, CK; ethylene, ET; gibberellins, GA; jasmonic acid; JA; salicylic acid, SA).

**Table 1.**
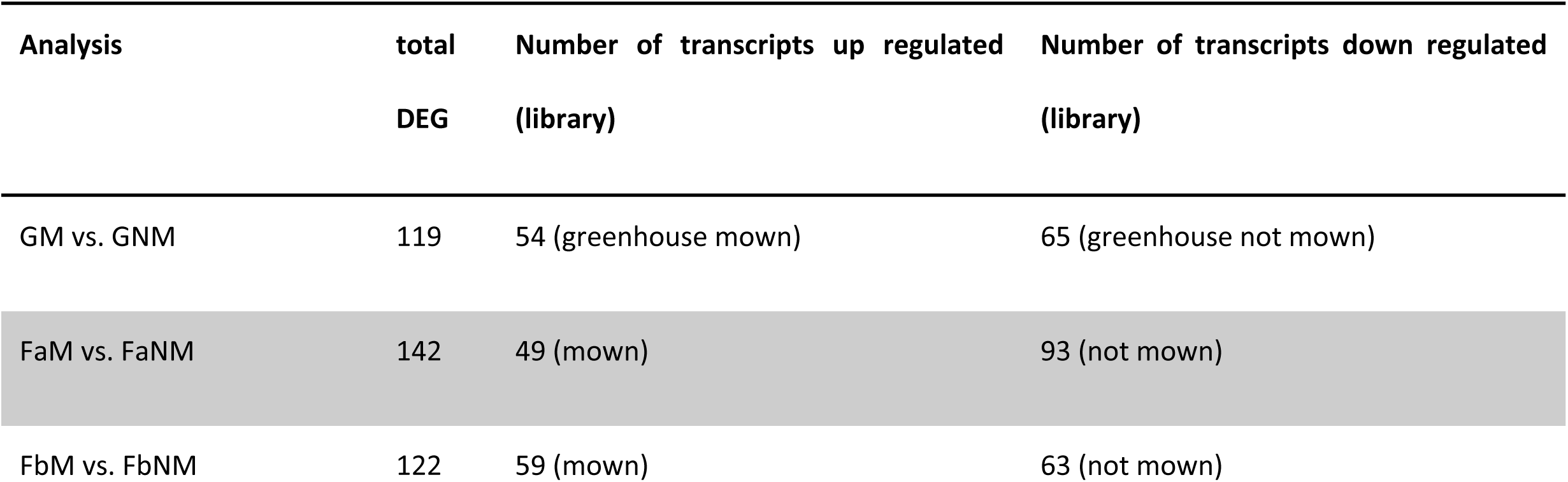
Table shows the numbers of differentially expressed transcripts (contigs) between libraries with changes above logfold 2. Up- or down regulation for each comparison is shown.

| Analysis | total<br>DEG | Number of transcripts up regulated<br>(library) | Number of transcripts down regulated<br>(library) |
| --- | --- | --- | --- |
| GM vs. GNM | 119 | 54 (greenhouse mown) | 65 (greenhouse not mown) |
| FaM vs. FaNM | 142 | 49 (mown) | 93 (not mown) |
| FbM vs. FbNM | 122 | 59 (mown) | 63 (not mown) |

RNA was extracted using NucleoSpin® RNA Plant Kit (Macherey-Nagel GmbH & Co. KG, Düren, Germany) according to the manufacturer’s instructions. Preparation of the cDNA libraries and the strand-specific sequencing was conducted by Eurofins Genomics (Ebersberg, Germany). The RNAs of four individuals were pooled for each RNA-Seq library and sequenced on an Illumina Hiseq2000 platform with chemistry v3.0, creating 2x 100 bp paired end reads.

In order to assess the effect of GA during the regrowth reaction of *T. pratense*, 14 red clover plants were mown as described in [25]. Of these plants, seven were used as control plants and seven plants were sprayed with 100 µM GA_3_ (Duchefa Biochemie B.V, Haarlem, The Netherlands) once per week as described in [37]. Different morphological characters (leaf number, length/width of leaflets, petiole length, number of inflorescences, and number of main shoots) were measured for four weeks.

### Assembly of reference transcriptome and annotation

The raw-read-quality of the RNA-Seq data was analyzed with FastQC (available online at: http://www.bioinformatics.babraham.ac.uk/projects/fastqc). Illumina adapter and low quality regions were trimmed using Trimmomatic [38] with ILLUMINACLIP, SLIDINGWINDOW:5:20 and MINLEN:50 options. Quality trimmed reads were pooled and digitally normalized [39]. Multiple d*e novo* assemblies were computed using Trinity [40] and Oases [41] with all odd k-mer parameters between 19 and 85. In addition, a genome guided assembly was performed using Trinity using the draft genome of *T. pratense* 1.0 (GCA_000583005.2) [12, 42]. The resulting contigs were screened for potential coding sequences (CDS) using TransDecoder (https://transdecoder.github.io/). The EvidentialGene pipeline (http://arthropods.eugenes.org/EvidentialGene/about/EvidentialGene_trassembly_pipe.html) was used to merge and filter the contigs based on the TransDecoder CDS prediction. Completeness of the final contig was confirmed by computing the mapping-rate of the non-normalized reads to the contigs. The raw sequence reads can be found at NCBI: PRJNA561285.

The contigs were uploaded to the “Sequence Analysis and Management System” (SAMS) [43] for functional annotation with the SwissProt [44], TrEMBL [45] and Phytozome [46] (e-value cutoff of 1e-5) databases. Additionally, attributes like gene name or functional description were extracted from the blast hits. Contigs were mapped to the *T. pratense* reference genome using gmap [47]. All non-Viridiplantae contigs were discarded. Transcription factors were identified using a blastp search of the protein sequences against the plant transcription factor database Potsdam (PlnTFDB) (48 [48], version 3.0, http://plntfdb.bio.uni-potsdam.de/v3.0/) protein database with an e-value cutoff of 1e-20. The files contain the functional annotation description of all transcripts e-Appendix (Table S11).

### Differential gene expression analysis, enrichment analysis, and classification of differentially expressed genes

Read counts for each contig of the final assembly in each sample were computed using RSEM [49] with bowtie mapping. To identify differentially expressed *T. pratense* genes (DEG) a pairwise comparison of all treatments was preformed using the DESeq2 [50] tool with FDR ≤ 0.01 and |logFoldChange| ≥ 2 between FaM and FaNM, FbM and FbNM; GM and GNM respectively. The top 20 DEG were determined for each comparison based on the expression strength (log2 fold change). Homologues in the next closest species and *A. thaliana* for each *T. pratense* candidate gene were searched based on the *T. pratense* genome sequence deposited in Phytozome [46]. TPM (transcript per million) values were calculated to estimate contig expression level (Wagner et al 2012).

We used the description and gene names obtained from TrEMBLE and SwissProt to search the UniProt [51], NCBI [52] and TAIR [53] databases to obtain further information (Table S8). Raw reads that were assembled to contigs, exhibiting a gene structure (ORF) and attained a putative annotation referred to below as genes.

### Blast2Go Analysis of *T. pratense* genomes

Two local BLAST searches [54] with word-size of 3, e-value of 1.0E-3 and HSP length cutoff of 33 were performed against the PlnTFDB using Blast2GO [55]. Only the blast hits with the highest similarity were used for further comparisons (number of BLAST hits = 1), sequences with similarity below 50% and an e-value higher than 1.0e-4 were omitted. The Blast2GO output was compared with an in-house python3-script utilizing NumPy (https://numpy.org/), Pandas (https://pandas.pydata.org/) and Seaborn (https://seaborn.pydata.org/) applying the list of transcription factors (TF) downloaded from PlnTFDB (http://plntfdb.bio.uni-potsdam.de/v3.0/) to the blast output and furthermore visualizing the generated datasets. We searched Uniprot database hits for development and phytohormone related genes. Subsequently, gene IDs of gibberellin genes we searched for matches within our annotated *T. pratense* transcriptomes. Matches were filtered based on TPM values and classified based on biosynthesis, catabolism activation/repression or signaling/response, corresponding expression patterns within the transcriptome have been identified additionally.

## Results

### RNA-Seq results, *de novo* assembly, and functional description of contigs

The RNA-Seq produced a total number of short reads between 44.7 and 58.1 million for each library with two exceptions (table S2) totaling 608,041,012 raw reads. The *de novo* assembly of the reference transcriptome of *T. pratense* produced 44,643 contigs, of which 41,505 contigs were annotated and 29,781 contigs were identified as plant specific. The minimum length of the contigs was 124 bp, the maximum length 1171.31 bp (Table S3). After the *de novo* assembly of the *T. pratense* transcriptome, each individual library was mapped back against the reference transcriptome individually, to determine the overall alignment rate, which was between 77.85 % and 90.32 % (Table S4).

**Table 2:**
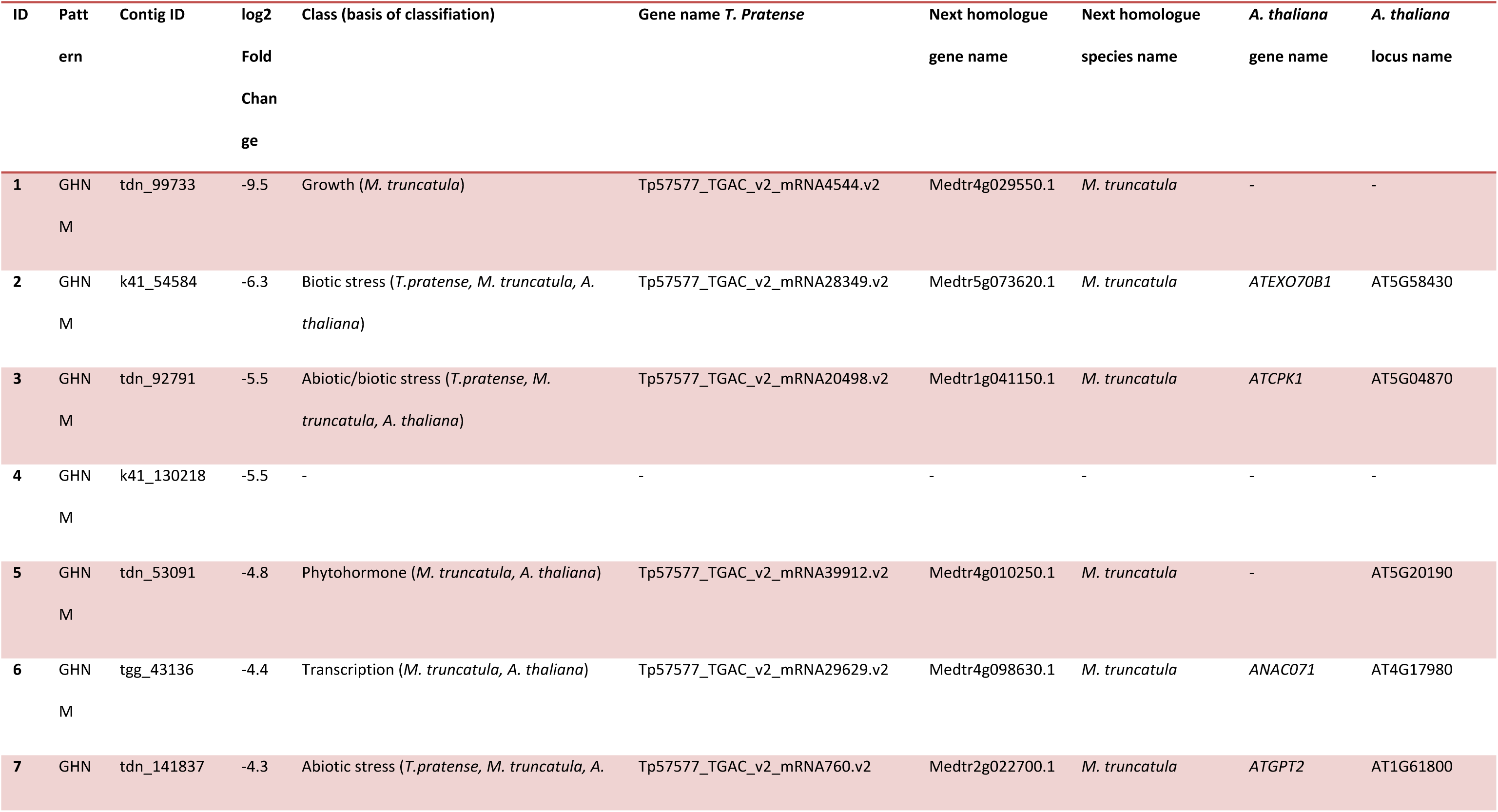
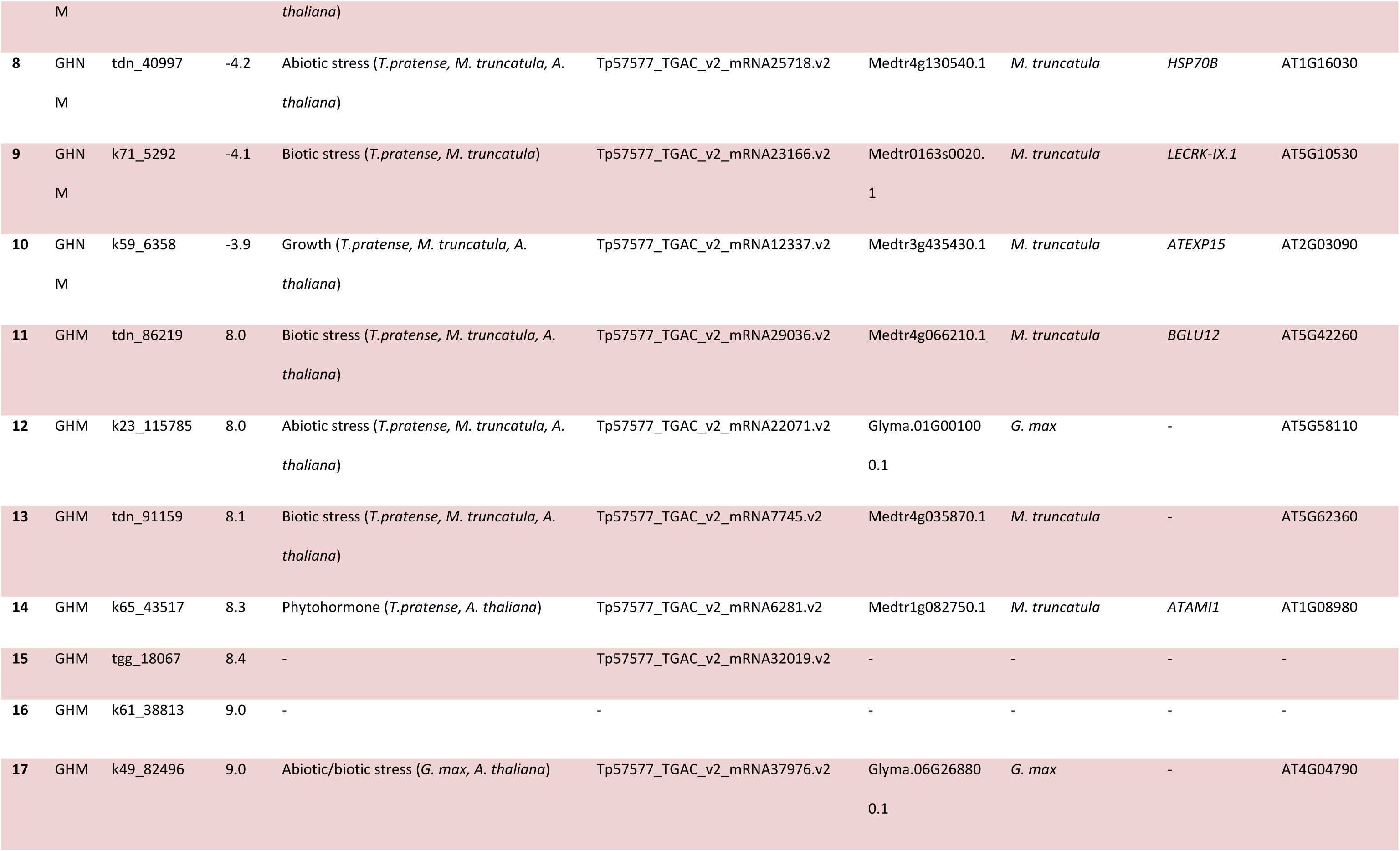
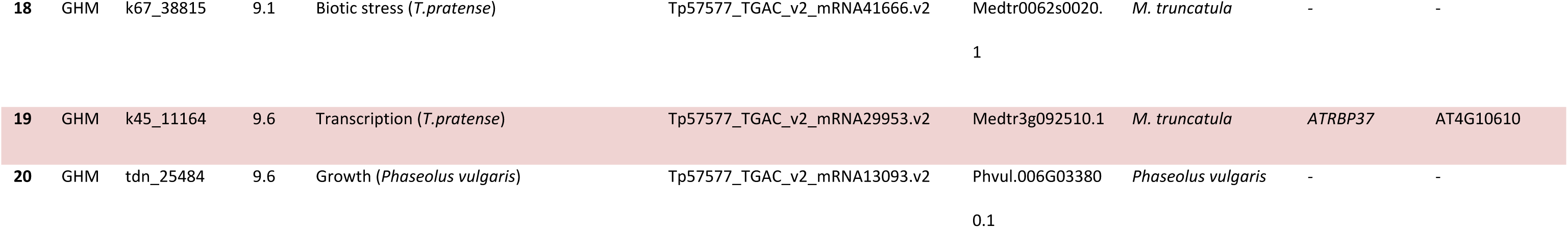
Top twenty differentially expressed genes of GM vs. GNM analysis. The table shows the transcript name, log2 fold change of the corresponding transcript, the library in which the transcript is upregulated (pattern), gene name based on *T. pratense* genome annotation, corresponding Phytozome description, gene name and species name of the next homologues and *A. thaliana* gene name and locus name based on information available on Tair.

**Table 3:**
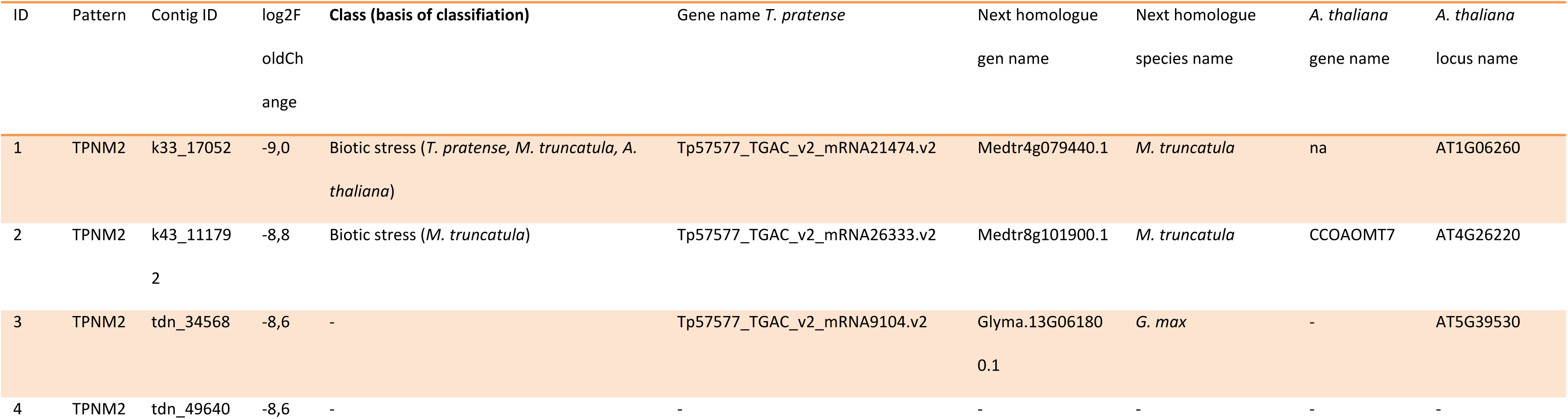
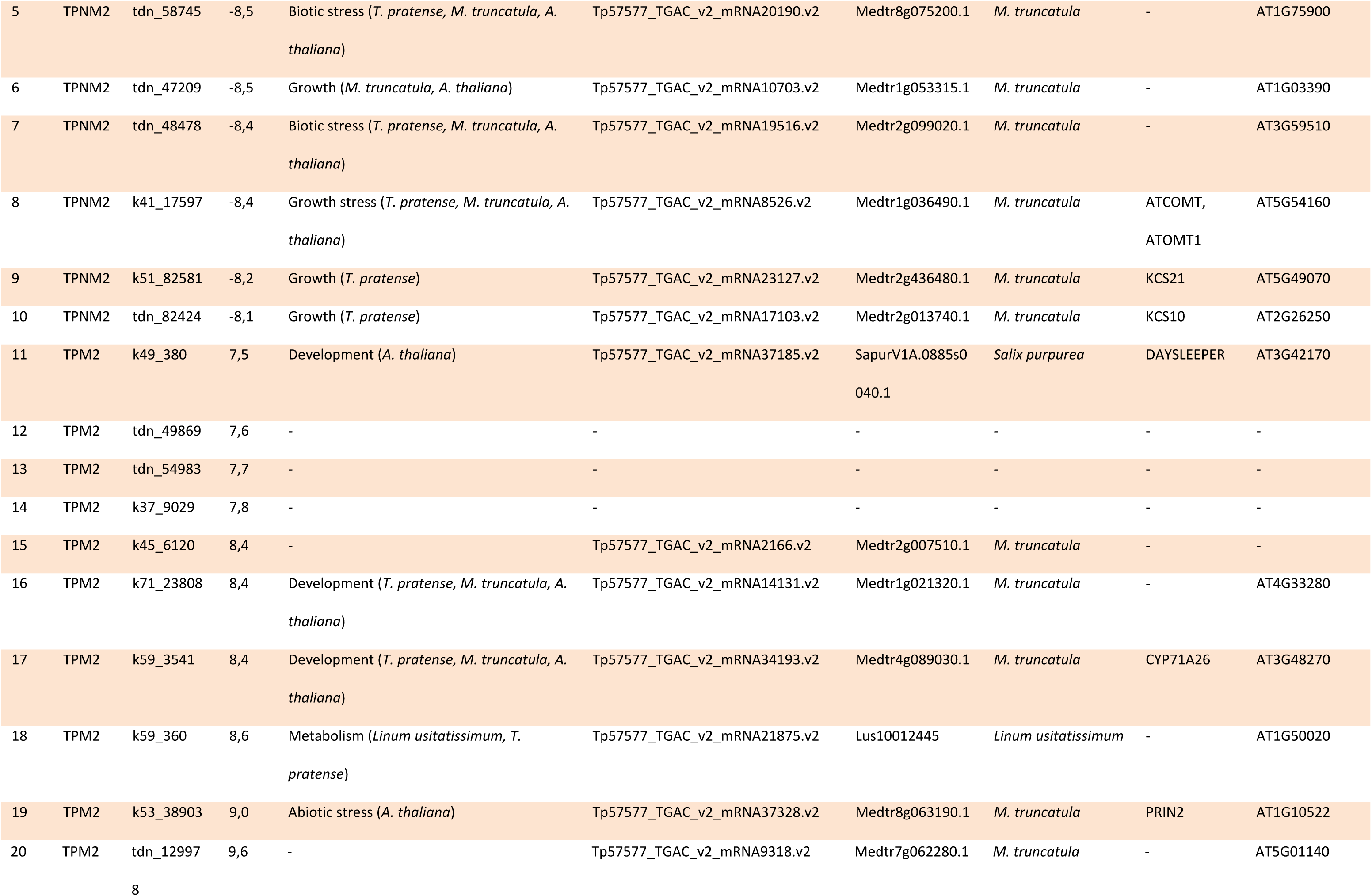
Top twenty differentially expressed genes of FaM vs. FaNM analysis. The table shows the transcript name, log2 fold change of the corresponding transcript, the library in which the transcript is upregulated (pattern), gene name based on *T. pratense* genome annotation, corresponding Phytozome description, gene name and species name of the next homologues and *A. thaliana* gene name and locus name based on information available on Tair.

**Table 4:**
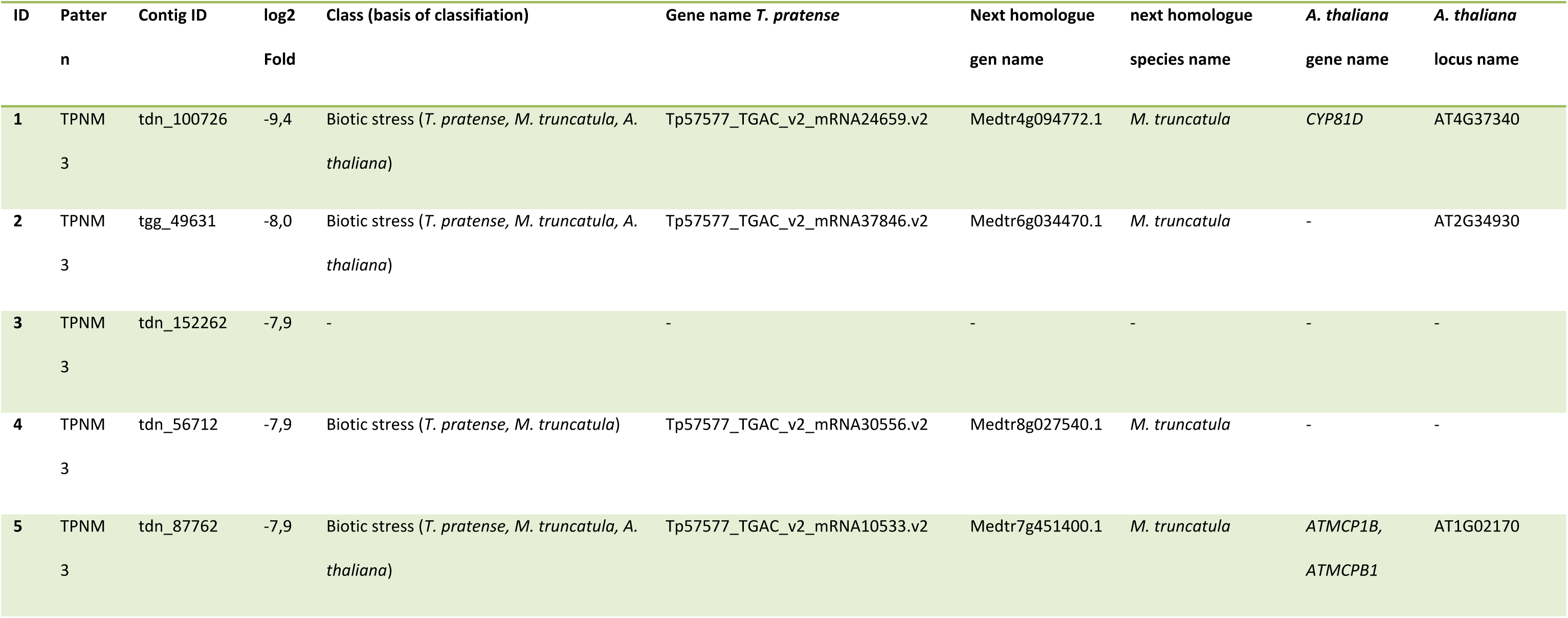
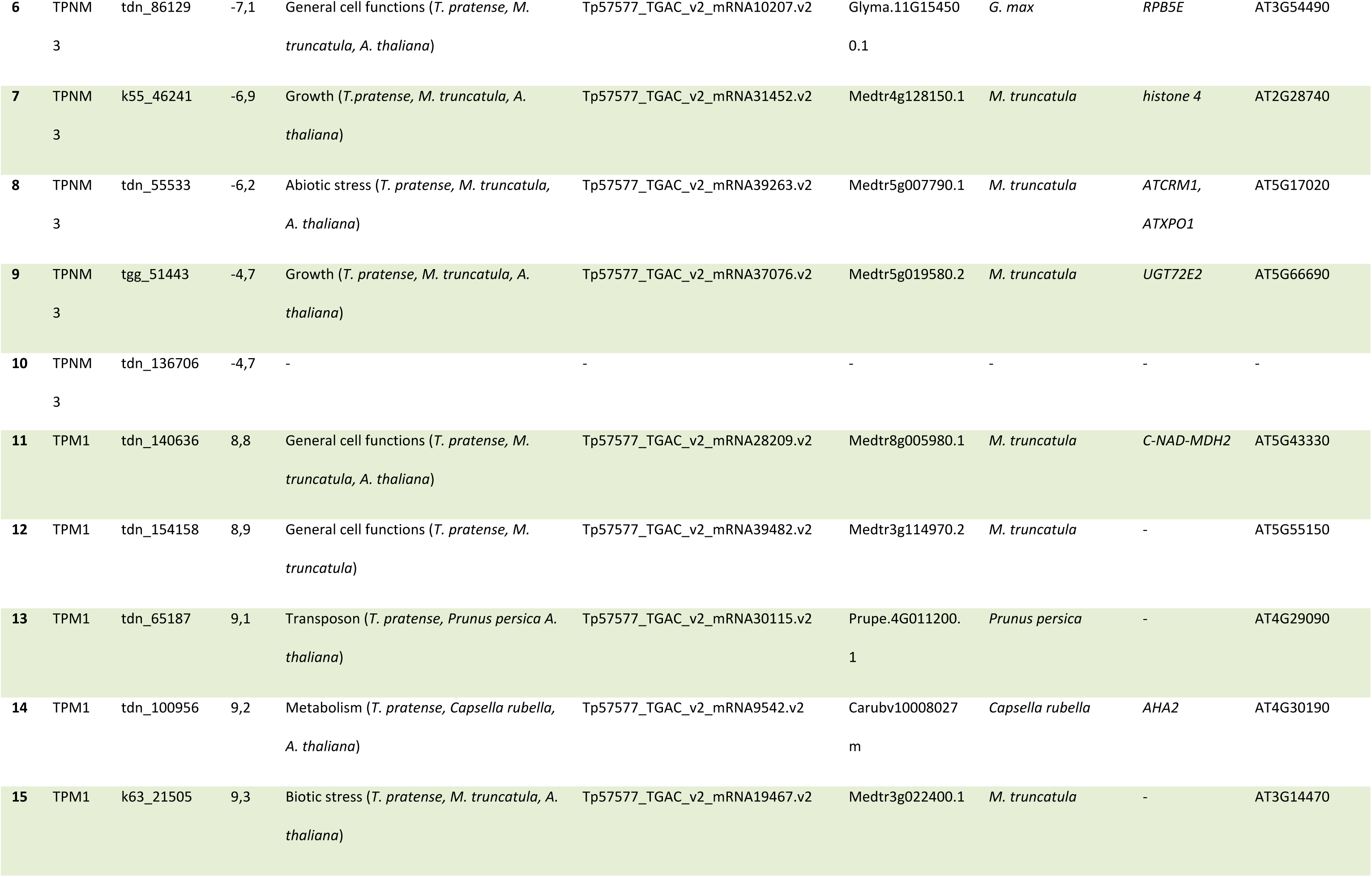
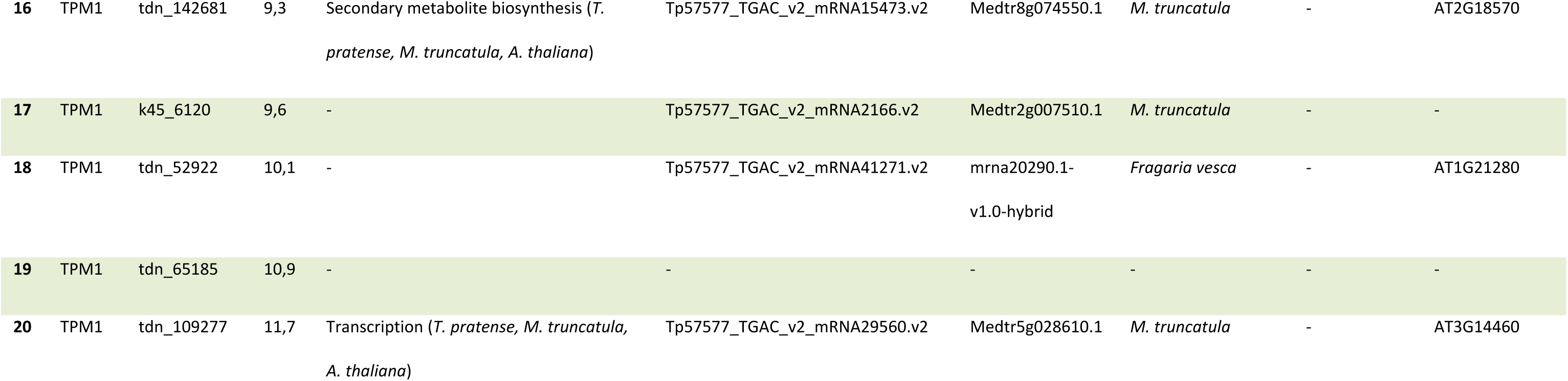
Top twenty differentially expressed genes of FbM vs. FbNM analysis. The table shows the transcript name, log2 fold change of the corresponding transcript, the library in which the transcript is upregulated (pattern), gene name based on *T. pratense* genome annotation, corresponding Phytozome description, gene name and species name of the next homologues and *A. thaliana* gene name and locus name based on information available on Tair.

63 % of the 44,643 contigs could be mapped to a known locus of the *T. pratense* genome annotation [12, 42]. 32 % could be mapped to an unknown locus of the *T. pratense* genome and 5 % could not be mapped to the *T. pratense* genome (Fig. S1). All plant-specific contigs were annotated with several databases (Table S5). To further verify the quality of our replicates, we identified the transcripts shared by the two replicates. We identified TPM values for each transcript and discarded transcripts with TPM values <1. Then we compared the transcripts of each library with each other and calculated the percentage of this number compared with the total number of transcripts within each library. The percentage of transcripts shared between the two replicates is between 90 % and 94 % for all treatments/localities, suggesting that the RNA-Seq data are reproducible (Table S6).

### Specific transcriptional regulator families are differentially expressed during the regrowth process

We were firstly interested to identify transcriptional regulators initiating and maintaining the regrowth morphology and mapped the transcriptome to the PlnTFDB to identify these transcriptional regulators. All members of a specific transcriptional regulator family (TRF) were *in silico* identified in the transcriptome and their expression was compared between mown and unmown plants (Fig. S2). Only those TRFs are shown for which at least 10% of the members showed significantly differential expression between mown and unmown conditions (Fig. 2).

**Fig. 2:**
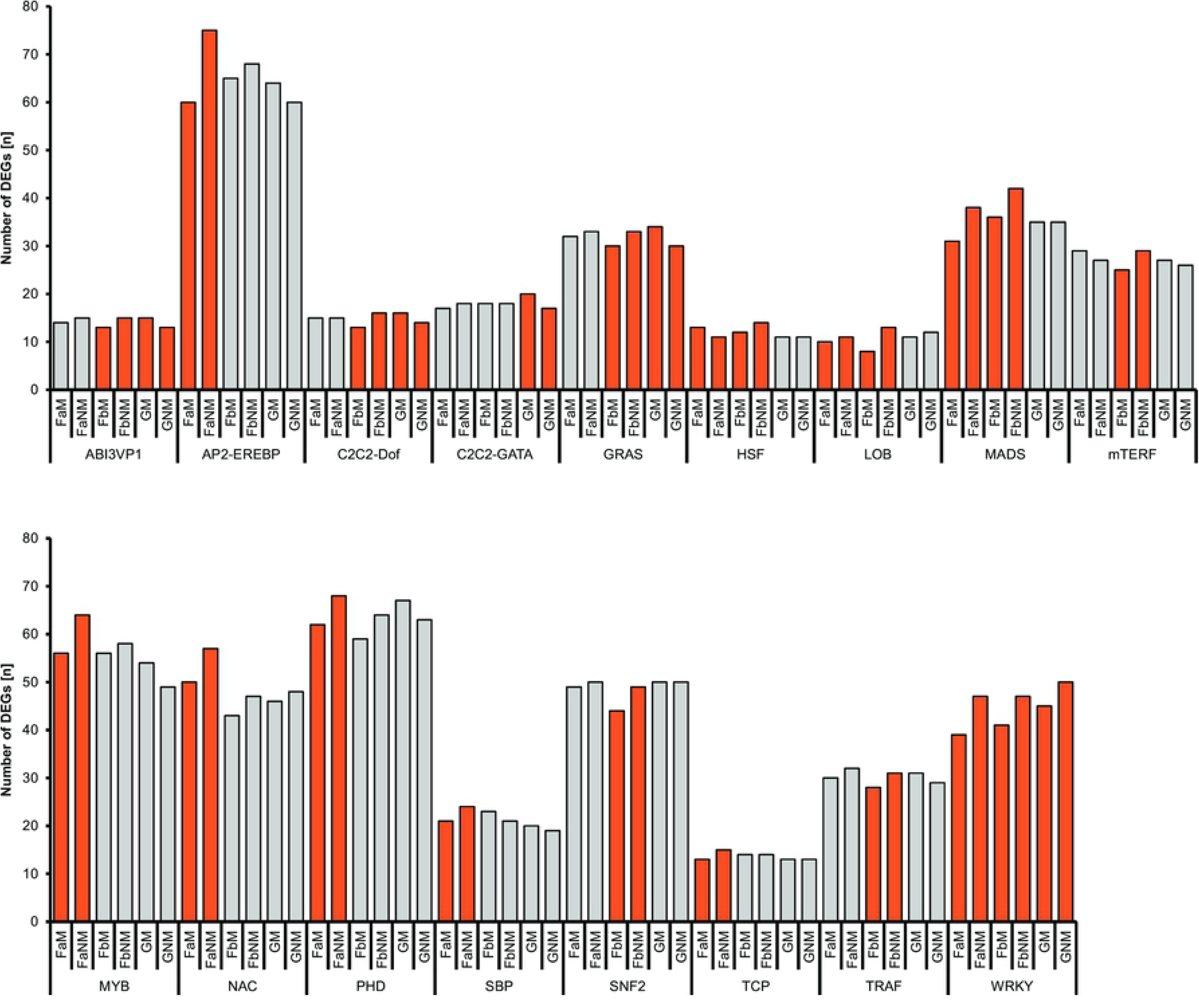
Differentially expressed TRF members in mown and not mown *T. pratense* plants. The y-axis shows the number of expressed contigs (TPM value over 5 TPM) that are members of the specific TRF. Names of the transcriptomes and TRFs are given on the x-axis. Expression of transcription factor members was compared in a pairwise manner (GM vs GNM, FaM vs FaNM, FbM vs FbNM). Shown are only those plant TRFs in which at least one of the comparisons resulted in a difference of more than 10% of the contigs significantly upregulated in either the mown or the unmown condition (orange bars).

17 TRFs were identified of which at least 10% of the members showed differential expression in the mown versus unmown comparisons (Fig. 2): ABI3VP1, AP2-EREBP, C_2_C_2_-Dof, C_2_C_2_-GATA, GRAS, HSF, LOB, MADS, mTERF, MYB, NAC, PHD, SBP, SNF2, TCP, TRAF, WRKY.

Two TRFs show expression activation upon mowing: a significant number of WRKY transcripts are up-regulated in mown plants regardless of the provenance. MADS-box transcripts were found upregulated as well, but only in the field-derived transcriptomes. Generally, only five of the 17 TRFs analyzed here showed significant changes in expression towards mowing in the greenhouse-derived plants suggesting that they react less strongly towards mowing than the field-derived plants. Six TRFs (AP2-EREBP, MYB, NAC, PHD, SBP, and TCP) show transcriptional changes in reaction to mowing only in field location a and three TRFs (mTERF, SNF2, TRAF) show this only in field location b suggesting that combination of biotic and abiotic factors with mowing differ between the two field locations.

Notably, only the C_2_C_2_-GATA TRF reacts towards mowing under greenhouse but not under field-conditions suggesting that transcriptional changes in reaction to other biotic and abiotic factors may overlay the regrowth reaction. Taken together, the TRF analysis shows that the reaction towards mowing induces transcriptional changes in only a subset of TRFs, suggesting that those play a major role in relieving the stress biomass loss and regrowth.

### Differentially expressed genes analysis reveals diverse subsets of genes involved in regrowth influenced by location and environmental conditions

To identify gene expression responses underlying the regrowth response after mowing digital gene expression analysis was performed comparing FaM vs FaNM; FbM vs FbNM; GM vs GNM to identify DETs (Table S12) from mown plants. Interestingly, using the log fold change 2, the number of DEG is rather similar in all comparisons, ranging from 119 (Gm vs. GNM) to 142 (FaM vs. FaNM) (Table 1).

We were then interested to identify developmental processes in greater detail that are required for the regrowth process. Thus, the results of the DEG analysis were restructured such that the DEG were grouped in 16 descriptive classes by database and literature mining (Table S7 and Table S8). Those classes describe major functional groups and serve to identify the potential role of a gene.

The results of the top 20 DEG showed that the greenhouse plants displayed more DEG involved in regrowth processes and less genes related to environmental conditions when compared with field plants. Most likely, the greenhouse grown plants displayed the regrowth reaction more prominently, as they grew under less stressful conditions than the field grown plants, for which more stress related DEG were observed (Fig. 1 B-D /Table 2-4).

Several functional groups show a similar pattern in the mown vs. unmown plants of all three locations: more genes related to biotic stress processes and metabolism were upregulated in the unmown locations (Fig. 1 B-D). In mown plants, more genes related to signaling and transposons were upregulated. Only a single functional group (growth) shows similar patterns in only the field locations suggesting that plants in the two field locations cope with very different habitat conditions and stress factors.

The photosynthesis- and phytohormone-related genes of field a show a similar pattern to the greenhouse plants as do the development- and signaling related genes. Genes related to development, general cell functions and transcription have similar patterns between field b and the greenhouse grown plants, such that more transcription - and development-related genes are upregulated in mown plants. DEG related to symbiosis were found upregulated in unmown plants grown in the greenhouse, even though these plants were fertilized. And unexpectedly, senescence-related genes are upregulated in mown plants of field A. However, because our analysis cannot discriminate between activating and repressing factors of senescence, we cannot conclude from our data if the mown plants have activated or repressed their senescence program.

The largest group of differentially expressed genes is the one related to biotic stress with up to 38% differentially expressed genes in one location (field b, Fig. 1 C). This suggests that different biotic stresses act upon the mown vs. unmown plants. A similar phenomenon can be observed for growth related processes, where up to 24% genes were upregulated in the mown and unmown plants indicating that different growth programs are active in mown vs. unmown plants.

Taken together we can state that mown plants in all locations change their regulatory programs upon mowing to cope with different biotic factors suggesting that they massively change their metabolism and signaling processes. Further, transposons are more active in mown plants. Apart from these conclusions, the molecular answer to substantial biomass loss differs between all three locations.

To find similarly regulated genes between the treatments and/or locations, a Venn diagram was generated to compare the number of shared significantly DEG within the “mown” samples and the “not mown” samples (Fig. 1 E-F, Table S9). Within the “mown” samples we detected no overlap between the groups with the exception of four genes that are differentially expressed and upregulated in “mown” condition and are shared between the two field transcriptomes (FbM and FaM (Fig. 1 E). Within the “not mown” samples also four genes are shared between the field transcriptomes (FbNM and FaNM)) and one is shared between the field b and the greenhouse (Fig. 1 F). No genes are shared between all three samples, neither in the “mown” treatment, nor in the “not mown” treatment. The genes that were shared between the transcriptomes belong to the main classes “growth”, “phytohormone”, “general cell functions”, “biotic stress”, “development” and “transcription” (Table S9).

Two of the genes could not be annotated. The annotated genes include for example genes tdn_60472 (shared between FaM/FbM, class: phytohormone), that was found to be the homolog of the *A. thaliana* locus AT1G75750, describing a GA-responsive GASA1 protein homolog. Another *A. thaliana* homolog was identified, Chitinase A (ATCHIA), shared between FaNM/FbNM (tdn_129843, class: biotic stress). In addition one gene was found, with a *T. pratense* annotation but no further description or homologs to *A. thaliana* (k45_6120, shared between FaM/FbM). This suggests that the molecular mechanisms directing regrowth overlaid by other processes, such as stress response which have a more dramatic impact on the number of DEG than growth processes have. The shared genes between the field conditions and the almost complete absence of shared genes between field and greenhouse indicates that the growth conditions in the field are more like each other, even when the fields are far apart from each other than any field to a greenhouse.

### Gibberellins are major players in the regrowth reaction

As phytohormones play a major role in the regulation of development and stress response, we identified DEGs related to phytohormone synthesis, homeostasis, transport, and signaling within all transcriptome comparisons (Table 1). DEG links for all major classes of phytohormones were identified, except for strigolactone. DEGs association to four phytohormones was most abundant: abscic acid (ABA, 8 DEGs), gibberellins (GA, 8 DEGs), salicylic acid (SA, 6 DEGs), and auxin (AUX, 5 DEGs) (Fig. 1 G). While ABA and SA are mainly involved in response to biotic and abiotic stresses, and AUX is known to play a major role in growth and development, we identified GA as a novel candidate phytohormone for regrowth response.

To learn more about the role of GA in the regrowth response, we identified 32 GA-related genes out of 151 within the transcriptomes of the greenhouse and the field grown plants, matching our selection criteria (TPM <5, involved in GA biosynthesis, signaling, GA responsive genes or catabolism, displaying certain expression patterns Fig. 3 A) and classified them according to their function in the GA biosynthesis and signaling processes (Table S13). Ranges of expression strength were calculated and color coded to compare expression patterns (Fig. 3 A).

**Fig. 3:**
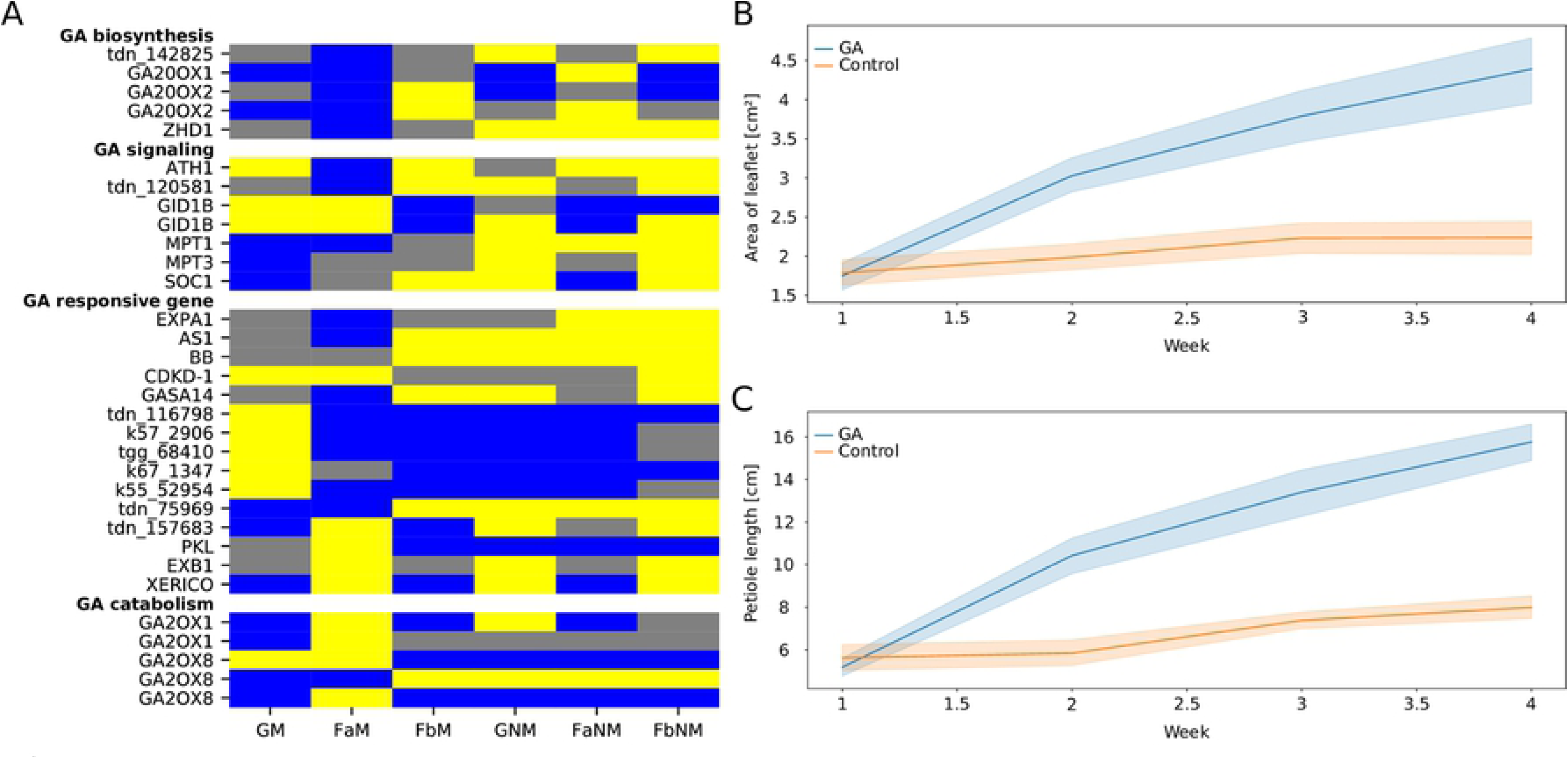
Analysis of GA related contigs and regrowth processes. A) Differentially expressed GA-related contigs within the *T. pratense* transcriptomes. Ranges of expression were calculated (0-39.99% blue (low expression), 40-59.99% grey (neutral expression), 60-100% yellow (high expression)) according to their TPM values. On the left, the gene names of the *T. pratense’s* closest *A. thaliana* homologs are given if available. B) and C) show morphological changes in leaves after GA treatment. B) leaflet area in cm², C) length of petioles in cm. The graphs show average values for each sampling date and 95% confidence interval. GA treated plants, blue; control plants, orange.

Five genes predicted to be involved in GA biosynthesis and signaling show similar expression differences between mown and unmown plants from at least two of the three locations: tdn_142825 (AT2G46590) and the homologs of *ZHD1, GID1B, MPT1*, and *MPT3*. Further, homologs of three GA responsive genes (tdn_75969 (*MYB 44*, AT5G67300), tdn_157683 (Os04g0670200, AT1G47128), and *XERICO* and homologs of two GA catabolism genes (*GA2OX1*, *GA2OX8*) react towards mowing. Interestingly, we also find many differences in expression between the field sites in the GA related genes suggesting fundamental differences in the living conditions between the two field sites that also impact regrowth after biomass loss.

When considering only the greenhouse grown plants, homologs of GA biosynthesis genes were not differentially regulated, but genes most likely involved in GA signaling such as *MPT1*, *MPT3*, and *SOC1* are down regulated in mown plants. Five homologs of GA responsive genes are upregulated in the mown plants, while three, among them *XERICO*, are down regulated. Further, of the three *GA2OXIDASE8* homologs encoded by the *T. pratense* genome, two are differentially expressed, one up- and the other down regulated upon mowing. Also, one *GA2OXIDASE1* homolog is down regulated in mown plants.

In summary, this suggests a highly dynamic response of several GA-related genes to mowing *or T. pratense*. Interestingly, we were unable to identify larger changes in the GA biosynthesis pathway of the greenhouse plants, but in GA catabolism genes, suggesting that GA availability in response to mowing is regulated by catabolism and signaling rather than by biosynthesis. Further, we identified two contrastingly regulated sets of genes acting in the mowing response.

### GA treatment after mowing induces specific changes to the regrowth response

Identification of several GA related genes that changed expression suggested an involvement of GA in the regulation of development after mowing. We were interested to corroborate this hypothesis experimentally and treated *T. pratense* plants with GA after mowing. Weekly GA application during the regrowth process led to significant and specific changes in morphology (Fig. 3 B, C). Previous work suggested that regrowing plants produce smaller and rounder leaflets with shorter petioles than uncut plants [25]. Number of leaves, shoots and inflorescences, leaf area and the roundness of leaflets were measured (Fig. 3 B, C, Suppl. Fig. 3). The first visible effects of GA treatment were recognized after 1.5 weeks, showing a significant difference in leaflet area between GA treated and control plants. Later it was observed that the petioles of treated plants were in average twice as long as petioles of untreated plants (16.7 ± 1.9 cm and 8 ± 1.2 cm, respectively). GA leaflets were with 4.7 ± 0.9 cm² almost double the size than those of untreated plants (2.4 ± 0.6 cm²). However, GA treated plants grew only 30% more total leaf area than control plants, because the control plants had more leaves than GA treated plants (Fig. S3 A, B, F, and G). Other morphological traits such as number of inflorescences, leaves, and shoots remained unaffected by the GA treatment. In summary, mown plants normally produce leaves with shorter petioles, restrict their leaflet area and their leaves become rounder. GA treatment partially alleviated these developmental changes such that the mown, GA treated plants produced larger leaves with longer petioles while the leaf shape was unaffected by GA treatment.

## Discussion

### RNA-Seq and assembly

The *de novo* assembly in combination with a reference-based approach for the annotation led to 44643 contigs of which 29781 could be annotated as plant-specific (Fig. S1). With the prior *de novo* assembly it was possible to attain 4051 additional contigs that could be not found within the genome of *T. pratense* 1.0 (GCA_000583005.2) [12, 42]. The estimated genome size of *T. pratense* is ∼440 Mbp [28]. The *T. pratense* transcriptome data in study was ∼55 Mbp in size, corresponding to ∼12.5% transcribed regions in the *T. pratense* genome, which is within the range of previously published transcriptomes (∼10% (42 Mbp) [56]). Interestingly, we found plant-specific, previously unreported contigs suggesting that the *T. pratense* genome might need improvement in terms of sequencing coverage and protein coding sequence annotation.

### Cell walls are remodeled after mowing

Our data analysis shows that several plant TRFs are predominantly involved in the regrowth reaction (Fig. 2 and S2). After massive biomass loss like mowing inflicts on *T. pratense*, plants firstly need to seal wounded tissues. Several transcriptional regulators are known to play a role in the tissue reunion processes were identified in *Solanum lycopersicum*, *Cucumis sativus*, and *A. thaliana* (reviewed in [36]). Homologs of these genes were also identified to be differentially regulated in the *T. pratense* transcriptome after mowing, such as several members of the Auxin Response Factor (ARF) family or the No Apical Meristem (NAM) family member *ANAC071*. [57] suggested that high levels of AUX induce the expression of *ANAC071* via ARF6 and ARF8 (in the upper part of incised stems), at the same time reduced AUX level directly after the cutting activate the expression of *RAP2.6L*. In addition auxin signaling via ARF6 and ARF8 influences JA synthesis, via the activation of *DAD1*, thus together with LOX2 increases *RAP2.6L* expression during tissue reunion in *A. thaliana* [57]. Further it was demonstrated that ANAC071 can as a transcription factor initiate the expression of members of xyloglucan endotransglucosylase/hydrolases family (XTH20 and XTH19) which recombine hemicellulose chains to drives the cell proliferation during tissue reunion [58].

Interestingly, we were able to identify all members of the cell wall remodeling pathway mentioned above, displaying distinct expression pattern with some of them upregulated in mown plants including for example *XTH32* (k69_7012, upregulated in FbM, tdn_94651, upregulated in GM, FaM and FbM), *XTH6* (tdn_91763, upregulated in GM), *XTH8* (k71_5058, upregulated in GM, FbM), *XTH9* (tdn_113578, upregulated in GM), *XTHA* (tdn_87930, upregulated in GM), *LOX2* (tdn_156279, upregulated FbM), and *ARF8* (tdn_156886 upregulated in GM, tdn_156890 upregulated in GM) (Table S10, S12 and S13). This is suggesting that the early steps in the regrowth reaction are conserved in core eudicots and that the cell wall remodeling processes continue at least two weeks after mowing.

### Biotic and abiotic stresses contribute to differential gene expression

RNA-Seq experiments create a large amount of raw data which requires significant downstream analysis to provide a biologically meaningful dataset. We thus compared those 20 genes, that differed most strongly in their expression between the different treatments and locations (see table 4-6 and Fig. 1 B-D). These comparisons revealed that the mown greenhouse plants show the highest percentage of genes possibly involved in regrowth processes. Contrasting, the field transcriptomes display patterns of abiotic and biotic stress reactions. Comparisons of the top 20 DEG of the unmown field transcriptomes showed that plants grown on field a and b face biotic stress more than abiotic stress. One of the upregulated genes in field a is a chitinase homolog suggesting that those plants are under attack of fungi and/or insects. Follow-up analyses to correlate environmental conditions, biotic and abiotic stresses monitored within the Biodiversity Exploratories with differential gene expression at the two field locations would be an interesting project but are beyond the scope of this work. In contrast, the top 20 DE transcripts of the greenhouse plants include phytohormone- and transcription-related genes, but also a high proportion of biotic and abiotic stress-related genes. This suggests that also these plants have to cope with stresses, but to a lesser extent. Thus, their regrowth reaction is more visible within the top 20 DEG. Generally, the non-mown plants show a much higher number of upregulated biotic stress-related genes during a phase in their life when senescence commences and they become more susceptible to pathogen attacks. The mown plants during their regrowth phase are not senescing and their younger organs seem be less affected by pathogens.

### GA related genes influence regrowth of *T. pratense*

GAs are involved in multiple aspects of plant development like cell elongation, flowering time regulation, and seed germination. Consequently, genes encoding for proteins involved in the synthesis, perception, and catabolism of the various GAs influence plant form. The RNA-Seq data presented showed a high abundance of GA associated genes (Fig. 1 G and 3) which might explain the morphological changes to mowing, such as rounder leaves, temporary dwarf-like appearance, and higher cumulative biomass production in mown plants [25]. Two of the genes, expressed higher in mown plants than in control clover plants are *GA20OX1* and *GA20OX2*. They are key enzymes of GA synthesis by producing precursors of the active GA forms (reviewed in [59]) and a deficiency in their activity is correlated with a dwarfed growth phenotype in *A. thaliana* and *O. sativa* [60, 61]. These mutants show also slow down cell division and expansion rate [62]. The upregulation of *GA20OX2* in mown plants may meet the enhanced demand of active GA to promote and sustain the regrowth to increase cell division and elongation. Similar to an increased GA synthesis, the expression of genes involved in reception of GA, such as the ortholog of AtGID1B is enhanced in mown plants. In *A. thaliana*, loss-of-function mutants of this GA receptor show a dwarfed phenotype [63] and over-expression of GID1 in *Medicago sativa* promotes biomass accumulation and leaf roundness [64].

In contrast to these GA responsive genes, GA2OX2 and GA2OX8 are involved in GA catabolism [65] and high levels of GA are known to activate the expression of degrading enzymes [66]. Both genes are expressed higher in not mown control plants than in mown plants and they inactivate the bioactive GAs GA2OX over-expression results in stunted plants and delayed flowering time [67, 68]. While mown clover plants show a higher expression of genes associated with growth activation, the ABA biosynthesis regulator XERICO is higher expressed in unmown plants (Table S13). It is a direct target of the DELLA protein RGL2 and in addition negatively regulated by GID1B [69, 70]. Thus, XERICO might restrict GA mediated growth to confer the drought adaptation of the not mown plants as more water is lost through the high leaf biomass.

In summary, mowing seems to trigger differential gene expression of GA activating enzymes and catabolic enzymes suggesting a dynamic GA response, but the gene expression patterns were not informative in respect to the consequences for the phenotype. When analyzing the morphological effects of GA application to mown plants (Fig. 3 A-C) we could show that external GA application lead to the disappearance of specific traits typical of the mowing response. Mown plants develop shorter petioles and produce a smaller leaf size area [25], but when treated with GA, leaves and petioles grow up the size seen in unmown plants.

The growth promoting abilities of GAs by cell expansion and proliferation via stimulating the degradation of growth-repressing DELLA proteins are well established [62]. The length increase of petioles in GA treated mown plants is in line with reported data from non-mown *Pisum sativum* (pea) plants, but in those, leaf sizes remained unchanged after GA treatment [71], suggesting a more specific role for GA in the regrowth reaction after biomass loss. Moreover, it was shown in *A. thaliana* previously, that elevated GA concentrations enhance cell division rates in the distal end of leaves (reviewed in [72]). If these results are transferred to *T. pratense* GA treatment should result in longer leaflets after GA treatment of mown plants. However, the leaf shape did not change, only the size increased suggesting a regrowth-specific shift of growth pattern which is unaffected by GA but similar to leaf shape of juvenile plants [25].

Interestingly, GA treatment of mown *T. pratense* plants does not generally lead to stronger longitudinal growth as leaves retained the round shape characteristic for untreated mown plants. These regrowth-specific characteristics can also be found in other species, for example in *A. thaliana*, *Fragaria ananassa*, *Duchesnea indica* and *G. max* GA treatment causes elongated petioles and increased leaf sizes and a more erect growth habit [73–76]. This proposes a new method to increase the accumulation of biomass, suitable for animal fodder. Previous experiments with the grasses *Leymus chinensis* and *Lolium perenne* showed GA action to be limited by N fertilization [77, 78]. Red clover, living in symbiosis with nitrogen fixing bacteria, is not dependent on additional N fertilization and can produce high-protein content biomass without fertilizer on poor soils.

## Acknowledgements

We thank Andrea Weisert for excellent technical help and Dietmar Haffer for skillfully raising the plants. We also thank Volker Wissemann and Birgit Gemeinholzer for continuous discussions on the project. We thank Hermann Finke for his help during visualization.

## Supplement Figures

Figure S1: Annotation Overview: A: Distribution of transcripts that could be mapped to the *T. pratense* genome, to a known locus and were annotated with *T. pratense* genome identifier. B: Distribution of transcripts that could be mapped to an unknown T*. pratense* gene locus. C: Distribution of transcripts that could not be mapped o the *T. pratense* genome. D: Distribution of transcripts of whole transcriptome representing all 12 libraries.

Figure S2 Differential expression of putative transcription factors of *T. pratense*. The Y axis denotes the number of expressed TF family members, the x axis shows the treatments and TF families. Orange bars indicate that >10% of the TF members are differentially expressed between the treatments, the red bars indicates that >5% are differentially expressed.

Figure S3: Plant architectural characteristics and growth habit of GA treated plants. A-E Measured, counted or calculated plant characteristics during phenotypic monitoring experiments. GA treated plants, blue; control plants, orange. Graphs show average values and 95% confidence intervals. Time is shown in weeks. Growth habit of control plants (left side) vs GA treated plants (right side), after approximately 2 weeks of GA treatment and regrowth (F), and after 4 weeks (G).

## References

1. Isobe S, Klimenko I, Ivashuta S, Gau M, Kozlov NN. First RFLP linkage map of red clover (Trifolium pratense L.) based on cDNA probes and its transferability to other red clover germplasm. Theor Appl Genet. 2003; 108: 105–112. doi: 10.1007/s00122-003-1412-z.

2. Isobe S, Sawai A, Yamaguchi H, Gau M, Uchiyama K. Breeding potential of the backcross progenies of a hybrid between Trifolium medium × T. pratense to T. pratense. Can. J. Plant Sci. 2002; 82: 395–399. doi: 10.4141/P01-034.

3. Eriksen J, Askegaard M, Søegaard K. Complementary effects of red clover inclusion in ryegrass-white clover swards for grazing and cutting. Grass Forage Sci. 2014; 69: 241–250. doi: 10.1111/gfs.12025.

4. Young ND, Debellé F, Oldroyd GED, Geurts R, Cannon SB, Udvardi MK, et al. The Medicago genome provides insight into the evolution of rhizobial symbioses. Nature. 2011; 480: 520–524. doi: 10.1038/nature10625.

5. Sato S, Nakamura Y, Kaneko T, Asamizu E, Kato T, Nakao M, et al. Genome Structure of the Legume, Lotus japonicus. Theor Appl Genet. 2008; 15: 227–239. doi: 10.1093/dnares/dsn008.

6. Schmutz J, Cannon SB, Schlueter J, Ma J, Mitros T, Nelson W, et al. Genome sequence of the palaeopolyploid soybean. Nature. 2010; 463: 178–183. doi: 10.1038/nature08670.

7. Schmutz J, McClean PE, Mamidi S, Wu GA, Cannon SB, Grimwood J, et al. A reference genome for common bean and genome-wide analysis of dual domestications. Nat Genet. 2014; 46: 707– 713. doi: 10.1038/ng.3008.

8. Varshney RK, Song C, Saxena RK, Azam S, Yu S, Sharpe AG, et al. Draft genome sequence of chickpea (Cicer arietinum) provides a resource for trait improvement. Nat Biotechnol. 2013; 31: 240–246. doi: 10.1038/nbt.2491.

9. Lonardi S, Muñoz-Amatriaín M, Liang Q, Shu S, Wanamaker SI, Lo S, et al. The genome of cowpea (Vigna unguiculata L. Walp.). Plant J. 2019; 98: 767–782. doi: 10.1111/tpj.14349.

10. Kaur P, Bayer PE, Milec Z, Vrána J, Yuan Y, Appels R, et al. An advanced reference genome of Trifolium subterraneum L. reveals genes related to agronomic performance. Plant Biotechnol J. 2017; 15: 1034–1046. doi: 10.1111/pbi.12697.

11. Dluhošová J, Ištvánek J, Nedělník J, Řepková J. Red Clover (Trifolium pratense) and Zigzag Clover (T. medium) - A Picture of Genomic Similarities and Differences. Front Plant Sci. 2018; 9: 724. doi: 10.3389/fpls.2018.00724.

12. Ištvánek J, Jaros M, Krenek A, Řepková J. Genome assembly and annotation for red clover (Trifolium pratense; Fabaceae). Am J Bot. 2014; 101: 327–337. doi: 10.3732/ajb.1300340.

13. Vega JJ de, Ayling S, Hegarty M, Kudrna D, Goicoechea JL, Ergon Å, et al. Red clover (Trifolium pratense L.) draft genome provides a platform for trait improvement. Sci Rep. 2015; 5: 17394. doi: 10.1038/srep17394.

14. Jahufer MZZ, Ford JL, Widdup KH, Harris C, Cousins G, Ayres JF, et al. Improving white clover for Australasia. Crop Pasture Sci. 2012; 63: 739. doi: 10.1071/CP12142.

15. Barrett BA, Faville MJ, Nichols SN, Simpson WR, Bryan GT, Conner AJ. Breaking through the feed barrier: options for improving forage genetics. Anim. Prod. Sci. 2015; 55: 883. doi: 10.1071/AN14833.

16. Řepková J, Nedělník J. Modern Methods for Genetic Improvement of Trifolium pratense. Czech Journal of Genetics & Plant Breeding. 2014: 92–99.

17. Řepková J, Nedělník J. Modern methods for genetic improvement of Trifolium pratense. Czech J. Genet. Plant Breed. 2014; 50: 92–99. doi: 10.17221/139/2013-CJGPB.

18. Dias PMB, Julier B, Sampoux J-P, Barre P, Dall’Agnol M. Genetic diversity in red clover (Trifolium pratense L.) revealed by morphological and microsatellite (SSR) markers. Euphytica. 2008; 160: 189–205. doi: 10.1007/s10681-007-9534-z.

19. Annicchiarico P, Proietti S. White clover selected for enhanced competitive ability widens the compatibility with grasses and favours the optimization of legume content and forage yield in mown clover-grass mixtures. Grass Forage Sci. 2010; 140: no-no. doi: 10.1111/j.1365-2494.2010.00749.x.

20. Ford JL, Barrett BA. Improving red clover persistence under grazing. Proceedings of the NZ Grassland Association. 2011; 73: 119–124.

21. Naydenova G, Hristova T, Aleksiev Y. Objectives and approaches in the breeding of perennial legumes for use in temporary pasturelands. Bio Anim Husb. 2013; 29: 233–250. doi: 10.2298/BAH1302233N.

22. Tiffin P. Mechanisms of tolerance to herbivore damage:what do we know. Evolutionary Ecology. 2000; 14: 523–536. doi: 10.1023/A:1010881317261.

23. Diaz S, Lavorel S, McIntyre SUE, Falczuk V, Casanoves F, Milchunas DG, et al. Plant trait responses to grazinga global synthesis. Global Change Biol. 2007; 13: 313–341. doi: 10.1111/j.1365-2486.2006.01288.x.

24. van Minnebruggen A, Roldán-Ruiz I, van Bockstaele E, Haesaert G, Cnops G. The relationship between architectural characteristics and regrowth in Trifolium pratense (red clover). Grass Forage Sci. 2015; 70: 507–518. doi: 10.1111/gfs.12138.

25. Herbert DB, Ekschmitt K, Wissemann V, Becker A. Cutting reduces variation in biomass production of forage crops and allows low-performers to catch up: A case study of Trifolium pratense L. (red clover). Plant Biol (Stuttg). 2018; 20: 465–473. doi: 10.1111/plb.12695.

26. Conaghan P, Casler MD. A theoretical and practical analysis of the optimum breeding system for perennial ryegrass. Irish Journal of Agricultural and Food Research. 2011; 50: 47–63.

27. Ortega F, Parra L, Quiroz A. Breeding red clover for improved persistence in Chile: a review. Crop Pasture Sci. 2014; 65: 1138. doi: 10.1071/CP13323.

28. Sato S, Isobe S, Asamizu E, Ohmido N, Kataoka R, Nakamura Y, et al. Comprehensive structural analysis of the genome of red clover (*Trifolium pratense* L.). DNA Res. 2005; 12: 301–364. doi: 10.1093/dnares/dsi018.

29. Shimizu-Sato S, Tanaka M, Mori H. Auxin-cytokinin interactions in the control of shoot branching. Plant Mol Biol. 2009; 69: 429–435. doi: 10.1007/s11103-008-9416-3.

30. Stafstrom J. Influence of Bud Position and Plant Ontogeny on the Morphology of Branch Shoots in Pea (Pisum sativum L. cv. Alaska). Annals of Botany. 1995; 76: 343–348. doi: 10.1006/anbo.1995.1106.

31. Briske DD, Richards JH. Plant responses to defoliation: a physiological, morphological and demographicevaluation. In: Bedunah DJ, Sosebee RE, editors. Wildland plants. Physiological ecology and developmental morphology. 1st ed. Denver, Colo.: Society for Range Management; 1995. pp. 635–710.

32. Kotova LM, Kotov AA, Kara AN. Changes in Phytohormone Status in Stems and Roots after Decapitation of Pea Seedlings. Russian Journal of Plant Physiology. 2004; 51: 107–111. doi: 10.1023/B:RUPP.0000011309.47328.23.

33. Li S, Strid Å. Anthocyanin accumulation and changes in CHS and PR-5 gene expression in Arabidopsis thaliana after removal of the inflorescence stem (decapitation). Plant Physiology and Biochemistry. 2005; 43: 521–525. doi: 10.1016/j.plaphy.2005.05.004.

34. Scholes DR, Wszalek AE, Paige KN. Regrowth patterns and rosette attributes contribute to the differential compensatory responses of Arabidopsis thaliana genotypes to apical damage. Plant Biol (Stuttg). 2016; 18: 239–248. doi: 10.1111/plb.12404.

35. Fischer M, Bossdorf O, Gockel S, Hänsel F, Hemp A, Hessenmöller D, et al. Implementing large-scale and long-term functional biodiversity research: The Biodiversity Exploratories. Basic and Applied Ecology. 2010; 11: 473–485. doi: 10.1016/j.baae.2010.07.009.

36. Asahina M, Satoh S. Molecular and physiological mechanisms regulating tissue reunion in incised plant tissues. J Plant Res. 2015; 128: 381–388. doi: 10.1007/s10265-015-0705-z.

37. Blazquez MA, Green R, Nilsson O, Sussman MR, Weigel D. Gibberellins promote flowering of arabidopsis by activating the LEAFY promoter. Plant Cell. 1998; 10: 791–800. doi: 10.1105/tpc.10.5.791.

38. Bolger AM, Lohse M, Usadel B. Trimmomatic: a flexible trimmer for Illumina sequence data. Bioinformatics. 2014; 30: 2114–2120. doi: 10.1093/bioinformatics/btu170.

39. Haas BJ, Papanicolaou A, Yassour M, Grabherr M, Blood PD, Bowden J, et al. De novo transcript sequence reconstruction from RNA-seq using the Trinity platform for reference generation and analysis. Nat Protoc. 2013; 8: 1494–1512. doi: 10.1038/nprot.2013.084.

40. Grabherr MG, Haas BJ, Yassour M, Levin JZ, Thompson DA, Amit I, et al. Full-length transcriptome assembly from RNA-Seq data without a reference genome. Nat Biotechnol. 2011; 29: 644–652. doi: 10.1038/nbt.1883.

41. Schulz MH, Zerbino DR, Vingron M, Birney E. Oases: robust de novo RNA-seq assembly across the dynamic range of expression levels. Bioinformatics. 2012; 28: 1086–1092. doi: 10.1093/bioinformatics/bts094.

42. Ištvánek J, Dluhošová J, Dluhoš P, Pátková L, Nedělník J, Řepková J. Gene Classification and Mining of Molecular Markers Useful in Red Clover (Trifolium pratense) Breeding. Front Plant Sci. 2017; 8. doi: 10.3389/fpls.2017.00367.

43. Bekel T, Henckel K, Küster H, Meyer F, Mittard Runte V, Neuweger H, et al. The Sequence Analysis and Management System – SAMS-2.0: Data management and sequence analysis adapted to changing requirements from traditional sanger sequencing to ultrafast sequencing technologies. Journal of Biotechnology. 2009; 140: 3–12. doi: 10.1016/j.jbiotec.2009.01.006.

44. Boutet E, Lieberherr D, Tognolli M, Schneider M, Bairoch A. UniProtKB/Swiss-Prot. Methods Mol Biol. 2007; 406: 89–112.

45. Bairoch A, Apweiler R. The SWISS-PROT protein sequence data bank and its supplement TrEMBL. Nucleic Acids Res. 1997; 25: 31–36. doi: 10.1093/nar/25.1.31.

46. Goodstein DM, Shu S, Howson R, Neupane R, Hayes RD, Fazo J, et al. Phytozome: a comparative platform for green plant genomics. Nucleic Acids Res. 2012; 40: D1178–86. doi: 10.1093/nar/gkr944.

47. Wu TD, Watanabe CK. GMAP: a genomic mapping and alignment program for mRNA and EST sequences. Bioinformatics. 2005; 21: 1859–1875. doi: 10.1093/bioinformatics/bti310.

48. Pérez-Rodríguez P, Riaño-Pachón DM, Corrêa LGG, Rensing SA, Kersten B, Mueller-Roeber B. PlnTFDB: updated content and new features of the plant transcription factor database. Nucleic Acids Res. 2010; 38: D822–7. doi: 10.1093/nar/gkp805.

49. Li B, Dewey CN. RSEM: accurate transcript quantification from RNA-Seq data with or without a reference genome. BMC Bioinformatics. 2011; 12: 323. doi: 10.1186/1471-2105-12-323.

50. Love MI, Huber W, Anders S. Moderated estimation of fold change and dispersion for RNA-seq data with DESeq2. Genome Biol. 2014; 15: 550. doi: 10.1186/s13059-014-0550-8.

51. The UniProt Consortium. UniProt: the universal protein knowledgebase. Nucleic Acids Res. 2016; 45: D158–69. doi: 10.1093/nar/gkw1099.

52. NCBI Resource Coordinators. Database resources of the National Center for Biotechnology Information. Nucleic Acids Res. 2016; 44: D7–19. doi: 10.1093/nar/gkv1290.

53. Berardini TZ, Reiser L, Li D, Mezheritsky Y, Muller R, Strait E, et al. The Arabidopsis Information Resource: Making and Mining the ‘Gold Standard’ Annotated Reference Plant Genome. Genesis. 2015; 53: 474–485. doi: 10.1002/dvg.22877.

54. Altschul SF, Gish W, Miller W, Myers EW, Lipman DJ. Basic local alignment search tool. Journal of Molecular Biology. 1990; 215: 403–410. doi: 10.1016/S0022-2836(05)80360-2.

55. Götz S, García-Gómez JM, Terol J, Williams TD, Nagaraj SH, Nueda MJ, et al. High-throughput functional annotation and data mining with the Blast2GO suite. Nucleic Acids Res. 2008; 36: 3420–3435. doi: 10.1093/nar/gkn176.

56. Yates SA, Swain MT, Hegarty MJ, Chernukin I, Lowe M, Allison GG, et al. De novo assembly of red clover transcriptome based on RNA-Seq data provides insight into drought response, gene discovery and marker identification. BMC Genomics. 2014; 15: 453. doi: 10.1186/1471-2164-15-453.

57. Pitaksaringkarn W, Ishiguro S, Asahina M, Satoh S. ARF6 and ARF8 contribute to tissue reunion in incised Arabidopsis inflorescence stems. Plant Biotechnology. 2014; 31: 49–53. doi: 10.5511/plantbiotechnology.13.1028b.

58. Pitaksaringkarn W, Matsuoka K, Asahina M, Miura K, Sage-Ono K, Ono M, et al. XTH20 and XTH19 regulated by ANAC071 under auxin flow are involved in cell proliferation in incised Arabidopsis inflorescence stems. Plant J. 2014; 80: 604–614. doi: 10.1111/tpj.12654.

59. Salazar-Cerezo S, Martínez-Montiel N, García-Sánchez J, Pérez-y-Terrón R, Martínez-Contreras RD. Gibberellin biosynthesis and metabolism: A convergent route for plants, fungi and bacteria. Microbiological Research. 2018; 208: 85–98. doi: 10.1016/j.micres.2018.01.010.

60. Rieu I, Ruiz-Rivero O, Fernandez-Garcia N, Griffiths J, Powers SJ, Gong F, et al. The gibberellin biosynthetic genes AtGA20ox1 and AtGA20ox2 act, partially redundantly, to promote growth and development throughout the Arabidopsis life cycle. Plant J. 2008; 53: 488–504. doi: 10.1111/j.1365-313X.2007.03356.x.

61. Spielmeyer W, Ellis MH, Chandler PM. Semidwarf (sd-1), “green revolution” rice, contains a defective gibberellin 20-oxidase gene. Proc Natl Acad Sci U S A. 2002; 99: 9043–9048. doi: 10.1073/pnas.132266399.

62. Achard P, Gusti A, Cheminant S, Alioua M, Dhondt S, Coppens F, et al. Gibberellin Signaling Controls Cell Proliferation Rate in Arabidopsis. Current Biology. 2009; 19: 1188–1193. doi: 10.1016/j.cub.2009.05.059.

63. Iuchi S, Suzuki H, Kim Y-C, Iuchi A, Kuromori T, Ueguchi-Tanaka M, et al. Multiple loss-of-function of Arabidopsis gibberellin receptor AtGID1s completely shuts down a gibberellin signal. Plant J. 2007; 50: 958–966. doi: 10.1111/j.1365-313X.2007.03098.x.

64. Wang X, Li J, Ban L, Wu Y, Wu X, Wang Y, et al. Functional characterization of a gibberellin receptor and its application in alfalfa biomass improvement. Sci Rep. 2017; 7: 41296. doi: 10.1038/srep41296.

65. Thomas SG, Phillips AL, Hedden P. Molecular cloning and functional expression of gibberellin 2-oxidases, multifunctional enzymes involved in gibberellin deactivation. Proc Natl Acad Sci U S A. 1999; 96: 4698–4703. doi: 10.1073/pnas.96.8.4698.

66. Yamaguchi S. Gibberellin metabolism and its regulation. Annu Rev Plant Biol. 2008; 59: 225– 251. doi: 10.1146/annurev.arplant.59.032607.092804.

67. Curtis IS, Hanada A, Yamaguchi S, Kamiya Y. Modification of plant architecture through the expression of GA 2-oxidase under the control of an estrogen inducible promoter in Arabidopsis thaliana L. Planta. 2005; 222: 957–967. doi: 10.1007/s00425-005-0037-7.

68. Sakamoto T, Kobayashi M, Itoh H, Tagiri A, Kayano T, Tanaka H, et al. Expression of a gibberellin 2-oxidase gene around the shoot apex is related to phase transition in rice. Plant Physiol. 2001; 125: 1508–1516. doi: 10.1104/pp.125.3.1508.

69. Ariizumi T, Hauvermale AL, Nelson SK, Hanada A, Yamaguchi S, Steber CM. Lifting della repression of Arabidopsis seed germination by nonproteolytic gibberellin signaling. Plant Physiol. 2013; 162: 2125–2139. doi: 10.1104/pp.113.219451.

70. Zentella R, Zhang Z-L, Park M, Thomas SG, Endo A, Murase K, et al. Global analysis of della direct targets in early gibberellin signaling in Arabidopsis. Plant Cell. 2007; 19: 3037–3057. doi: 10.1105/tpc.107.054999.

71. DeMason DA, Chetty VJ. Interactions between GA, auxin, and UNI expression controlling shoot ontogeny, leaf morphogenesis, and auxin response in Pisum sativum (Fabaceae): or how the uni-tac mutant is rescued. Am J Bot. 2011; 98: 775–791. doi: 10.3732/ajb.1000358.

72. Nelissen H, Gonzalez N, Inzé D. Leaf growth in dicots and monocots: so different yet so alike. Curr Opin Plant Biol. 2016; 33: 72–76. doi: 10.1016/j.pbi.2016.06.009.

73. Guttridge CG, Thombson PA. The Effect of Gibberellins on Growth and Flowering of Fragaria and Duchesnea. J Exp Bot. 1964; 15: 631–646. doi: 10.1093/jxb/15.3.631.

74. Leite VM, Rosolem CA, Rodrigues JD. Gibberellin and cytokinin effects on soybean growth. Sci. agric. (Piracicaba, Braz.). 2003; 60: 537–541. doi: 10.1590/S0103-90162003000300019.

75. Tsukaya H, Kozuka T, Kim G-T. Genetic control of petiole length in Arabidopsis thaliana. Plant Cell Physiol. 2002; 43: 1221–1228. doi: 10.1093/pcp/pcf147.

76. Hisamatsu T, King RW, Helliwell CA, Koshioka M. The involvement of gibberellin 20-oxidase genes in phytochrome-regulated petiole elongation of Arabidopsis. Plant Physiol. 2005; 138: 1106–1116. doi: 10.1104/pp.104.059055.

77. Cai Y, Shao L, Li X, Liu G, Chen S. Gibberellin stimulates regrowth after defoliation of sheepgrass (Leymus chinensis) by regulating expression of fructan-related genes. J Plant Res. 2016; 129: 935–944. doi: 10.1007/s10265-016-0832-1.

78. Morvan-Bertrand A, Ernstsen A, Lindgard B, Koshioka M, Le Saos J, Boucaud J, et al. Endogenous gibberellins in Lolium perenne and influence of defoliation on their contents in elongating leaf bases and in leaf sheaths. Physiol Plant. 2001; 111: 225–231. doi: 10.1034/j.1399-3054.2001.1110214.x.

